# Characterization of Gene Expression Changes Over Healthy Term Pregnancies

**DOI:** 10.1101/229773

**Authors:** Anna K. Knight, Anne L. Dunlop, Varun Kilaru, Dawayland Cobb, Elizabeth J. Corwin, Karen N. Conneely, Alicia K. Smith

**Affiliations:** Genetics and Molecular Biology Program, Emory University, Atlanta, GA; Emory University School of Nursing, Atlanta, GA; Department of Gynecology and Obstetrics, Emory University School of Medicine, Atlanta, GA; Department of Human Genetics, Emory University School of Medicine, Atlanta, GA; Department of Psychiatry, Emory University School of Medicine, Atlanta, GA

**Keywords:** RNA, transcript, PBMC, maternal, pregnancy complication, hemoglobin, trimester

## Abstract

During pregnancy, women experience numerous physiological changes but, to date, there is limited published data that characterize accompanying changes in gene expression over pregnancy. This study sought to characterize the complexity of gene expression over the course of pregnancy among women with healthy pregnancies. Subjects provided a venous blood sample during early (6-15 weeks) and late (22-33 weeks) pregnancy, which was used to isolate peripheral blood mononuclear cells prior to RNA extraction. Gene expression was examined for 63 women with uncomplicated, term deliveries. We evaluated the association between weeks gestation at sample collection and expression of each transcript. Of the 16,311 transcripts evaluated, 439 changed over pregnancy after a Bonferroni correction to account for multiple comparisons. Genes whose expression changed over pregnancy were associated with oxygen transport, the immune system, and host response to bacteria. Characterization of changes in gene expression over the course of healthy term pregnancies may enable the identification of genes whose expression predicts complications or adverse outcomes of pregnancy.

## Introduction

Pregnancy is characterized by extensive physiological changes including increased blood volume, elevated levels of estrogen and progesterone, changes in metabolism, and shifts in the maternal immune system in order to accommodate the demands of the growing fetus[1, 2]. Despite these carefully regulated physiological changes that occur throughout the 40 weeks of an average pregnancy, no study has yet described accompanying changes in maternal blood gene expression among healthy women with full term, uncomplicated pregnancies.

To date, studies of gene expression focus on comparisons of pregnant women who are healthy to those with autoimmune disorders. This has been prompted, in part, by the observation that women with some autoimmune disorders report alleviation of their symptoms during pregnancy [3]. For example, a recent study of peripheral blood from 20 women with rheumatoid arthritis and 5 healthy controls identified 4,710 genes that differed in their expression over pregnancy and the postpartum period. The genes identified were enriched for a variety of pathways including immune pathways, signal transduction, and disease-related pathways. Interestingly, several of the genes identified by this study, are members of the alpha defensin family and are involved in immune and defense responses [4]. Another study identified 1,286 transcripts whose expression levels changed over three timepoints in pregnancy and the postpartum period for women with rheumatoid arthritis and controls. These transcripts were enriched for a variety of pathways including hematopoietic cell lineage and toll-like receptor signaling. The authors concluded that the identified pathways may contribute to immunomodulation and thus a reduction in rheumatoid arthritis symptoms during pregnancy with these changes in gene expression being largely reversed postpartum [5]. Gilli and colleagues sampled multiple sclerosis patients and healthy controls before pregnancy, during each trimester, and post-partum to identify 404 transcripts whose expression differed between multiple sclerosis patients and healthy controls. A refined signature of 347 transcripts was then used to evaluate gene expression in patients and controls in the ninth month of pregnancy, at which time the signature could no longer distinguish patients from controls [6]. These studies indicate that gene expression does change over pregnancy, potentially leading to alleviation of some of the symptoms of autoimmune disorders.

Although such studies have been informative for identifying pathways that change in women with autoimmune disorders during pregnancy, if the groups are not sampled at comparable stages of gestation, the findings may be more difficult to interpret or replicate in independent studies. This study sought to characterize gene expression changes over pregnancy in a cohort of healthy women with uncomplicated term deliveries. The pathways identified provide further insight into the connection between gene expression and physiological changes that occur over the course of a healthy, full term pregnancy, and will serve as a resource for ongoing studies of pregnancy complications and adverse pregnancy outcomes.

## Results

### Study Subjects

63 pregnant African American women provided venous blood samples between 6-15 weeks and again between 22-33 weeks gestation, representing the late first/early second trimesters and the late second/early third trimesters. The clinical and demographic features of these women are summarized in Table 1. At the time of enrollment, 7 women (12%) report using tobacco within the last 30 days, and 4 women (7%) report drinking alcohol within the last 30 days. Despite this, no woman delivered a neonate that was small for gestational age or experienced any complications over their pregnancies (see Methods for additional details).

**Table 1:** Clinical and demographic characteristics of the 63 women included in the study.

| | Mean ± SD |
| --- | --- |
| Maternal Age (years) | 25.4 ± 4.5 |
| Parity | 1.2 ± 1.1 |
| Gravidity | 2.8 ± 1.6 |
| Length of Gestation (weeks) | 39.4 ± 1.0 |
| Birthweight (grams) | 3318 ± 411.8 |
|  | N (%) |
| Delivery Type |  |
| Vaginal | 55 (87) |
| C-section | 8 (13) |
| Insurance Type |  |
| Medicaid | 46 (73) |
| Private | 17 (37) |
| Education |  |
| Some High School | 11 (17) |
| High School Graduate | 17 (27) |
| Some College | 25 (40) |
| College Graduate | 7 (11) |
| Graduate School | 3(5) |

### Changes in cell proportions over pregnancy

Studies have reported changes in a range of cell types and immune activation profiles over pregnancy[7-9]. Consistent with those previous reports, we observed changes in the proportion of cell types in maternal PBMCs over the course of pregnancy. There was a significant increase in the proportion of monocytes (p=.001), along with a significant decrease in the proportion of B cells (p=.03), and natural killer cells (p=.004) based on week of gestation (Fig 1). Estimates of cell composition used for this study are not comprehensive, and there may be additional changes in cell composition or function that are not currently captured. To account for such changes in cell type and the potential for other known (i.e. age) or unknown confounders, we used surrogate variable analysis (SVA), which identifies and estimates sources of expression heterogeneity to increase power to detect true and replicable associations. Post-hoc evaluation of surrogate variables suggests that they reflect changes in cell composition, age, and control for technical variables such as batch.

**Figure 1:**
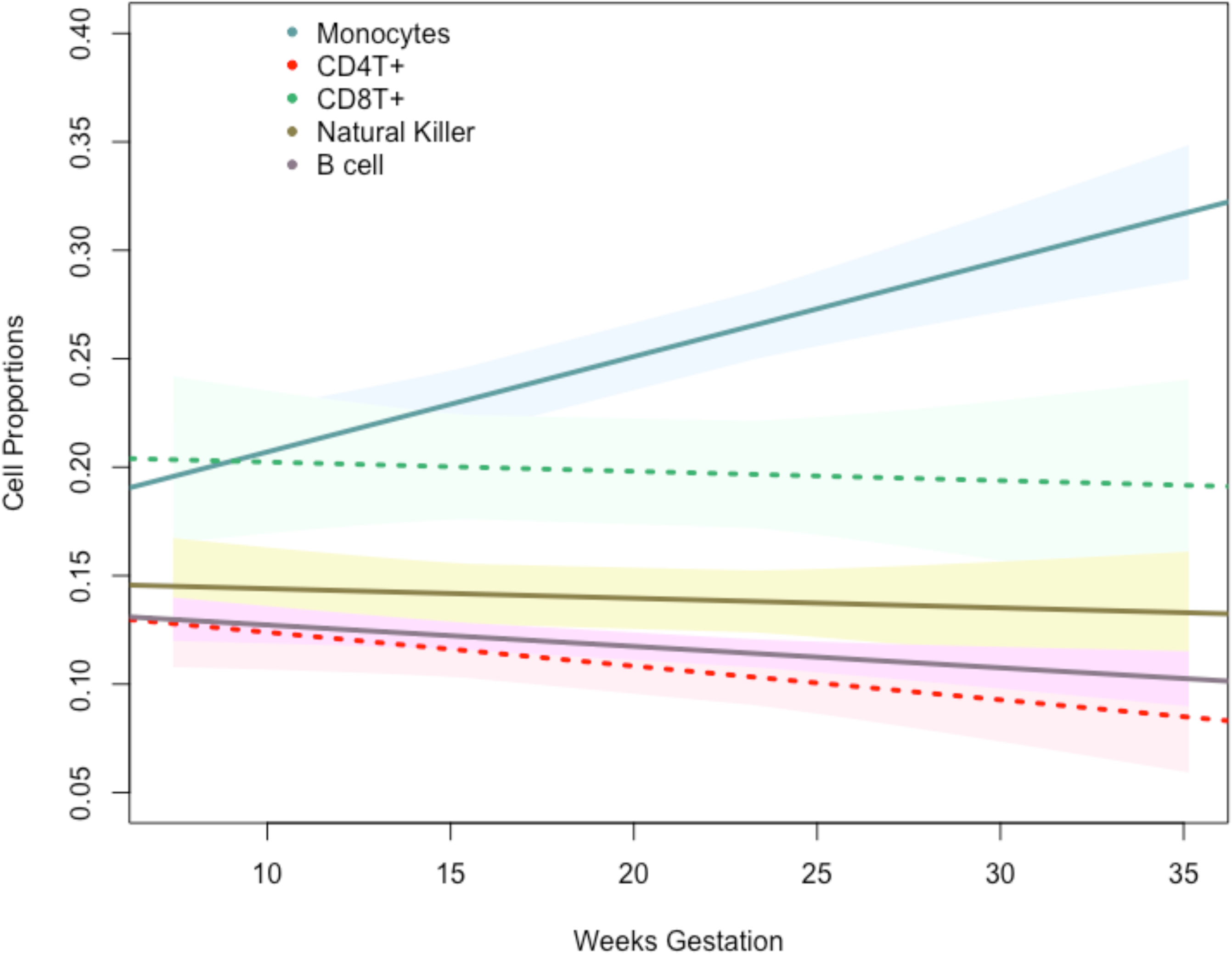
Cellular composition changes over pregnancy. Monocytes increase over pregnancy, whereas B cells and natural killer cells decrease over pregnancy. Other evaluated cell types do not change. Regression lines are shown with 95% confidence bands. Solid line indicates significance (p<.05). The x-axis represents the weeks of gestation at sample collection and the y-axis represents cell proportions.

### Gene expression changes over pregnancy

Of the 16,311 transcripts that were evaluated in this genome-wide analysis (Fig 2), 439 associated with weeks of gestation at sample collection, after adjusting for multiple testing (Table S1). The majority (69.6%) of these genes increased in expression over the course of pregnancy. Genes whose expression changed were enriched for multiple biological processes (Table 2). Of note, many of these genes are in involved in oxygen transport or the hemoglobin complex (e.g. *AHSP*, *HBD*, *HBM*, and *HBQ1*). In addition to increased expression of genes involved in oxygen transport, other associated biological processes emphasize changes in the immune system known to occur over pregnancy. Several of these immune processes are associated with response to microbes, including the antibacterial humoral response and the innate immune response in mucosa (Table 2). Key genes in these pathways are members of the alpha defensin family, including *DEFA1*, *DEFA4*, and *DEFA1B*(Fig 3).

**Figure 2:**
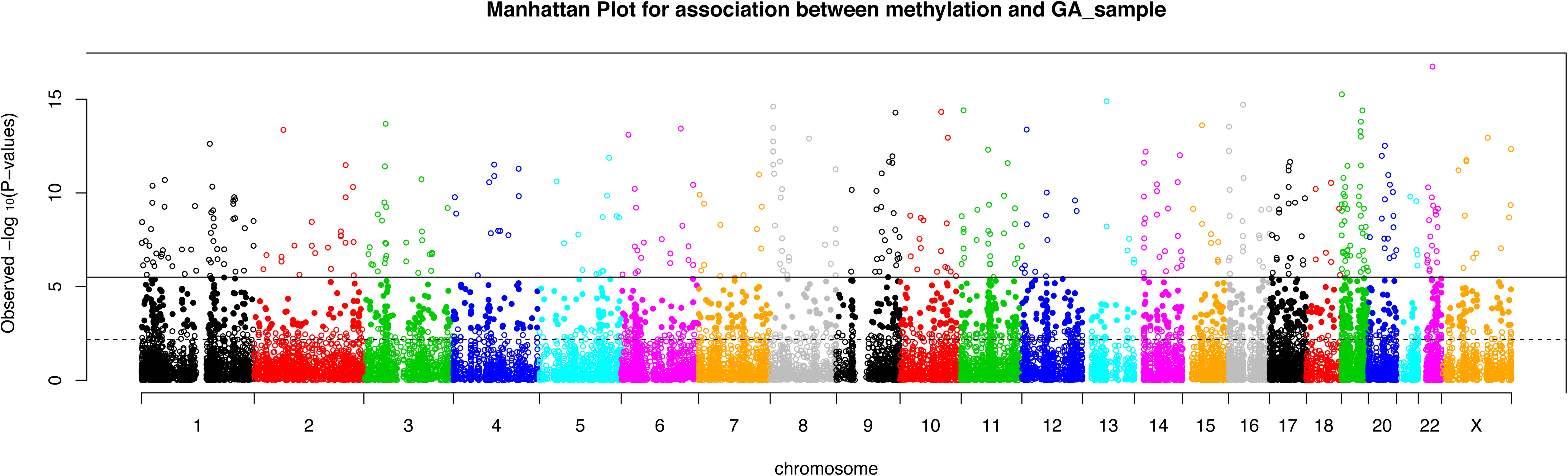
Genome-wide distribution of transcripts that change over pregnancy. Manhattan plot depicting associations with gene transcripts that change over pregnancy. Expression probes are oriented by position for each chromosome (x-axis). The y-axis represents the significance level for each test. The solid line denotes experiment-wide significance based on a Bonferroni adjustment, and the dotted horizontal lines denotes experiment-wide significance based on a Benjamini-Hochberg adjustment.

**Table 2:** Biological Processes associated with genes whose expression changes over pregnancy.

| Term | GO Identifier | Count | p-value* |
| --- | --- | --- | --- |
| oxygen transport | GO:0015671 | 6 | 0.0074 |
| defense response to fungus | GO:0050832 | 7 | 0.0081 |
| antibacterial humoral response | GO:0019731 | 8 | 0.012 |
| leukocyte migration | GO:0050900 | 13 | 0.012 |
| killing of cells of other organism | GO:0031640 | 5 | 0.041 |
| innate immune response in mucosa | GO:0002227 | 6 | 0.044 |

\* p-values presented after a Benjamini-Hochberg correction for multiple testing.

**Figure 3:**
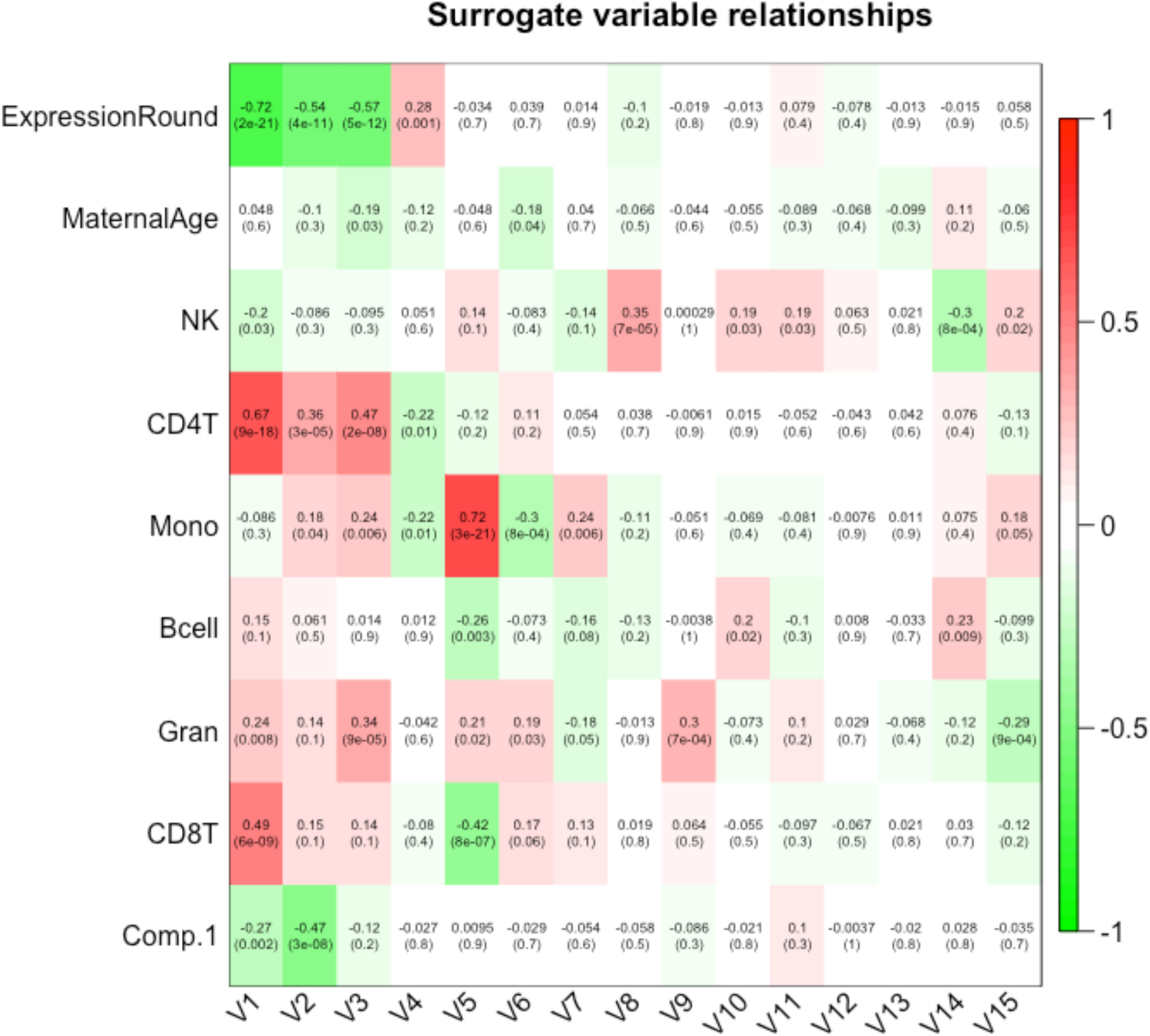
Genes in the alpha defensing family that change over pregnancy. The x-axis represents weeks gestation at sample collection, and the y-axis represents the log2-transfomed expression levels for each transcript. a) *DEFA4* (ILMN_1753347; p=2.45x10^-15^) b) *CEACAM8*(ILMN_1806056; p=1.57x10^-14^) c) *DEFA1* (ILMN_1679357; p=3.37x10^-14^) d) *DEFA1B* (ILMN_2102721; p=1.79x10^-13^).

In contextualizing our study findings with the existing literature, we attempted to compare the genes whose expression levels change in healthy pregnancies to those that have been reported to change in women with autoimmune disorders. In some cases, complete gene lists were not provided [5, 6], limiting our ability to make direct comparisons. For example, Weix and colleagues reported 19 selected candidate genes that differed in pregnancy and differed to a greater degree in pregnant women with rheumatoid arthritis. Of those candidate genes, only 1 *(STAT1)* differed in our genome-wide analysis. The study by Mittal and colleagues reported the change in expression of 256 genes over pregnancy in women with rheumatoid arthritis and controls [4], 179 of which were evaluated in our study. Of these, 55.3% change over pregnancy in our cohort of healthy women. This is a substantially higher number than would be expected by chance (99/179; p<2.2x10^-16^).

## Discussion

In this study, we identified 439 transcripts that were associated with weeks gestation at sample collection in uncomplicated term pregnancies. These transcripts were enriched for several key pathways that provide insight into the mechanisms underlying the physiological changes necessary to support a healthy pregnancy.

A widely accepted phenomenon in pregnancy is a change in the maternal immune system. In general, pregnancy is associated with a shift from a pro-inflammatory state in the first trimester to an anti-inflammatory state in the second trimester, with renewed inflammation during the third trimester and at parturition [2]. The pro-inflammatory state in the first trimester is likely a result of implantation and placentation with the anti-inflammtory state of the second trimester being a period of rapid fetal growth and development during which a more symbiotic relationship between mother, fetus, and placenta exists. Finally in the third trimester, renewed inflammation leads to the processes which can initiate labor and delivery. Consistent with this and other studies, our results suggest that monocytes increase over the course of pregnancy, while B cell and natural killer cell proportions decrease [10, 11]. This shift has been further explored in studies of autoimmune disorders and pregnancy, with several pathways involved being identified that may contribute to reductions, or in other cases, increases in autoimmune symptoms during pregnancy [3-6]. However, such studies often do not control for differences in cellular proportions, increasing the chance that they will identify genes whose expression is not entirely attributable to the disorder of interest or are otherwise difficult to interpret. Thus, it is vital for future studies to estimate and adjust for cellular heterogeneity, as cell type changes over pregnancy.

In this longitudinal analysis of genome-wide changes in expression across gestational week, we identified gene expression changes that supports functional differences in the immune system during pregnancy. Such immune system changes must be carefully regulated so that the mother is protected from bacterial and viral infection without negatively impacting the body’s tolerance of the fetus. However, the immune systems of pregnant women are less able to appropriately respond to several types of bacterial infections and related complications, including *Listeria monocytogenes* and *Neisseria gonorrhoeae* [12-14]. Bacterial infections have been associated with preterm birth and spontaneous abortion [15-17]. Evidence of host-microbe interactions in pregnancy is demonstrated in both the gene-level results and through enrichment of genes involved in antibacterial humoral response and innate immunity in the gene ontology analysis. Specific genes related to bacterial response include several alpha defensin genes. These genes encode antimicrobial peptides (AMPs) that have been associated with microbicidal activity and host defense, and are present in the female reproductive tract during pregnancy [18, 19]. Understanding changes in the immune system and host-microbe interactions during pregnancy may provide future insight into how the immune system acts to protect both the mother and the fetus thus allowing us to more effectively treat and prevent potential maternal and fetal effects of systemic infection.

This study also identified genes involved in oxygen transport that change over pregnancy, which support the physiological changes identified in previous studies [20-23]. Some of the most pronounced adaptations in pregnancy are related to increased blood volume, higher rates of erythropoiesis, and increased coagulation. Maternal physiology tends toward a more hypercoagulable state during pregnancy, likely in an effort to limit delivery complications such as post-partum hemorrhage. Previous studies have linked abnormal coagulation, specifically within the placental complex, with preeclampsia, demonstrating the importance of regulation of coagulation for healthy pregnancies[24, 25].

Erythropoiesis increases over the course of pregnancy, potentially to accomadate larger blood volumes and prepare for the acute loss of this volume which occurs at the time of delivery. However, this study evaluated PBMCs, which do not include mature erythrocytes[26]. Still, recent studies have reported the expression of hemoglobin in non-erythroid cells, such as macrophages, epithelial, mesangial, cervical, and endometrial cells, and have also reported functions of hemoglobin other than oxygen transport including antioxidant defense, nitrite reduction, and reactive oxygen species scavenging [27, 28]. We hypothesize that the hemoglobin-associated genes identified in this study reflect non-canonical expression in response to the physiological strain that pregnancy places on the body, potentially resulting in further demands for such alternative functions.

This is the first study to comprehensively characterize changes in gene expression longitudinally over the course of uncomplicated pregnancies. This cohort has been prospectively recruited and characterized, resulting is an invaluable asset for studying pregnancy progression [29]. However, this study does have limitations. Though it is the largest longitudinal study of gene expression changes in pregnancy, it still has a modest sample size. A larger group of subjects is likely to identify more genes that change over the course of pregnancy. Despite this, we were able to identify hundreds of genes whose expression change in PBMCs after accounting for confounding factors and multiple testing. However, we cannot extrapolate these changes to other cell types that may be relevant for pregnancy. This study also uses array-based technology, which does not allow for characterization of novel or alternatively spliced transcripts that may play vary during pregnancy. Finally, we did not have access to samples spanning the entirety of pregnancy, including before 8 weeks and after 33 weeks gestation.

Despite these limitations, we show that gene expression changes significantly over the course of pregnancy, especially in pathways related to oxygen transcript and response to microbes. Our results provide a framework for future studies on changes in gene expression over pregnancy. Knowledge of genes associated with normal changes may potentially allow for identification of abnormal patterns of gene expression associated with pregnancy and delivery complications that could be tested as potential biomarkers. Future studies should examine gene expression over pregnancy in the context of pregnancy complications and other pre-existing conditions, while acknowledging that very subtle changes in the timing of sample collection may confound the results or complicate their interpretation.

## Methods

### Study Subjects

Pregnant African American women were recruited, enrolled, and underwent data collection as part of the Emory University African American Vaginal, Oral, and Gut Microbiome in Pregnancy Cohort Study, as described previously [29]. To summarize, women were recruited from March 2014 through August 2016 at outpatient prenatal care clinics affiliated with two Atlanta metro area hospitals, Emory University Midtown Hospital and Grady Memorial Hospital. Women were eligible for inclusion if they self-identified as African American, were between 18-40 years of age, had a singleton pregnancy, had less than four previous births, and were able to understand written and spoken English. Additional exclusion criteria for this analysis included the following indicators of high-risk pregnancy or pregnancy complications: gestational diabetes, hypertension, intrauterine growth restriction, preterm birth, preeclampsia, eclampsia, hemolysis, elevated liver enzymes, low platelet count (HELLP) syndrome, hyperemesis gravidarum, oligohydramnios, chorioamnionitis, macrosomia, preterm premature rupture of membranes (pPROM), or fetal intolerance of labor. Fetal death before labor and congenital abnormalities of the fetus were criteria for post-enrollment exclusion. Demographic data were collected through self-report questionnaires. Clinical obstetrical data (including estimated due date, gestational age at delivery, pregnancy complications, labor and delivery course) were ascertained via abstraction of the prenatal and labor and delivery medical chart by a qualified physician. This study was approved by the Emory University Institutional Review Board.

#### Biological Sample Collection

63 women contributed two samples each over the course of their pregnancy. The first sample was collected at 6-15 weeks, and the second sample was collected at 22-33 weeks. During each prenatal visit, an additional 12 mL of venous blood was drawn using the same needle stick as for the routine blood draws. PBMCs were isolated from whole blood using a Ficoll density gradient and were stored in AllProtect Buffer (Qiagen) at -80 °C until a simultaneous DNA and RNA extraction using the AllPrep RNA/DNA Mini Kit (Qiagen) was performed according to manufacturer’s instructions. DNA quantification and quality was assessed using the Quant-it Pico Green kit (Invitrogen). RNA quantification and quality was assessed using the Aligent RNA Nano 6000 Kit and Bioanalyzer 2100.

#### RNA Expression Analysis

For each subject, gene expression was assessed for ~47,000 transcripts using the HumanHT-12 v4 BeadChip (Illumina). Briefly, 750 ng of RNA was directly hybridized to the BeadChip according to manufacturer’s instructions. The BeadChips were scanned using the iScan scanner, and the raw data was analyzed using the Expression Module of GenomeStudio Software (Illumina). Two samples with detection p values >.01 for more than 90% of probes were excluded. Probes detected in less than 10% of samples were also excluded; 16,311 probes passed quality control. Data was then quantile-normalized and log2 transformed prior to association testing. RNA expression data can be accessed through NCBI’s Gene Expression Omnibus, GEOXXXX.

#### Cell type composition estimation

DNA methylation was interrogated for each subject using either the HumanMethylation450 or MethylationEPIC BeadChip, which measures methylated and unmethylated signal for >450,000 and >850,000 CpG sites across the genome, respectively. Initial data quality control was performed using the R package CpGassoc [30]. Any CpG site with low signal or missing data for greater than 10% of samples was removed, and any sample with missing data for greater than 5% of CpG sites was removed. Cross-reactive probes were removed [31]. Following quality control, 449,094 probes were included in subsequent analyses. Beta values (β) were calculated for each CpG site as the ratio of methylated (M) to methylated and unmethylated (U) signal: β=M/M+U. Beta-mixture quantile normalization was performed as previously described [32]. Cell type proportions (CD8T+, CD4T+, natural killer, B cell, monocytes, and granulocytes) were estimated as previously described from DNA methylation data [33]. Associations between cell type proportions and gestational age were examined using a linear-mixed effects model. DNA methylation data can be accessed through NCBI’s Gene Expression Omnibus, GEOXXXX.

#### Genome-wide Expression Analysis

We used linear mixed-effects modeling implemented in the R package “nlme” to interrogate associations between gene expression and weeks gestation at sample collection[34].The R package sva was used to estimate surrogate variables to control for potentially confounding factors, including cell type [35]. Surrogate variable analysis was used instead of a covariate adjustment as only a few cellular subtypes are well defined using current methods, and changes in the composition of unmeasured cell types may have a large influence during pregnancy. The 15 significant surrogate variables were included as covariates in the model. Figure S1 shows the associations between each surrogate variable and common confounders. A random effects term was included in the model to account for repeated sampling of the same person. A Bonferroni correction was applied to account for multiple testing. Pathway analysis for associated genes was performed using DAVID [36].

## Acknowledgements

We are grateful to the participants enrolled in this cohort who provided samples and medical record data for this study. We are grateful to Misti B. Patel for providing insights into the clinical relevance of this work.

---

**Table S1**: CpG sites that change over pregnancy.

**Figure S1:** Association of surrogate variables and common confounders. R (p-value).

